# The number and pattern of viral genomic reassortments are not necessarily identifiable from segment trees

**DOI:** 10.1101/2023.09.20.558687

**Authors:** Qianying Lin, Emma E. Goldberg, Thomas Leitner, Carmen Molina-París, Aaron A. King, Ethan O. Romero-Severson

## Abstract

Reassortment is an evolutionary process common in viruses with segmented genomes. These viruses can swap whole genomic segments during cellular co-infection, giving rise to novel progeny formed from the mixture of parental segments. Because large-scale genome rearrangements have the potential to generate new phenotypes, reassortment is important to both evolutionary biology and public health research. However, statistical inference of the pattern of reassortment events from phylogenetic data is exceptionally difficult, potentially involving inference of general graphs in which individual segment trees are embedded. In this paper, we argue that, in general, the number and pattern of reassortment events are not identifiable from segment trees alone, even with theoretically ideal data. We call this fact the fundamental problem of reassortment, which we illustrate using the concept of the ‘first-infection tree’, a typically but not always counterfactual genealogy that would have been observed in the segment trees had no reassortment occurred. Further, we illustrate four additional problems that can arise logically in the inference of reassortment events and show, using simulated data, that these problems are not rare and can potentially distort our perception of reassortment even in small data sets. Finally, we discuss how existing methods can be augmented or adapted to account for not only the fundamental problem of reassortment but also the four additional situations that can complicate the inference of reassortment.

## Introduction

Reassortment is the evolutionary process by which viruses with segmented genomes exchange genetic material [1, 2]. When a cell becomes dually infected, that is, infected by two distinct viruses, the resulting viral offspring can contain a mixture of genomic segments from each parent [3]. The genetic diversity and fitness of segmented viruses such as influenza A virus [4, 5], rotaviruses [6, 7], and bunyaviruses [8, 9] are influenced by reassortment. For example, in bunyaviruses, the M segment encodes the viral glycoprotein that mediates both cell fusion and immune responses [10]. By swapping M segments, therefore, the reassortant can lead to much more severe illness in humans compared to its parents. As a concrete example, the hemorrhagic Ngari virus emerged from reassortment between the only mildly pathogenic Bunyamwera and Batai viruses [11, 12]. Other impacts of reassortment include escaping vaccine-elicited immunity [13], increasing fitness in humans [14], and getting access to new hosts [15]. Understanding the molecular mechanism, the evolutionary process, and the epidemiological implications of reassortment is essential for anticipating and combating emerging infectious diseases [16], improving public health strategies [17], and guiding the development of novel diagnostics, therapeutics, and vaccines [5]. Further, reassortment can be thought of as a special case of recombination, one in which the breakpoints are fixed and correspond to the endpoints of each segment [3].

In population genetics, a phylogenetic tree is commonly used to represent evolutionary history over time, in which the root represents the most recent common ancestor (MRCA) of all the sampled genomes, and internal nodes represent a splitting of one lineage into two, denoting the ancestral relationships between sampled tips [18]. For viral pathogens, this pattern of branching reflects both the epidemiology of who infected whom, and also exactly who within the population was sampled [19]. The logic underlying reconstructions of reassortment is that all else being equal, topological discordance between segment trees is caused by historical reassortment events (Fig. 1). For example, the observation that two viruses that co-cluster in a tree inferred from one genomic segment, but are quite distant in a tree inferred from another segment, is likely the result of a recent reassortment event. This logic is the foundation of existing reassortment inference methods, which follow one of two distinct approaches. One approach begins with segment trees and then attempts to synthesize their topological incongruence into an ancestral reassortment graph (ARG), wherein specific reassortment events correspond to reticulations, *i*.*e*., partial merging of two ancestor lineages [20– 23]. This is analogous to the practice of reconciling gene trees to infer recombination [24, 25]. These methods are intuitively appealing but they have trouble pinpointing the timing of reassortment events in dated segment trees [26]. A second approach seeks to infer an ARG—or in the case of Ref. [27], a very closely related data structure called a phylogenetic network—jointly with the segment trees using the likelihood under a coalescent-with-reassortment model [27, 28]. These approaches are generally based on an extension of the Kingman coalescent [29, 30], that obtains a high level of computational efficiency by only considering the events in the population that are specifically ancestral to the sampled genomes. In the coalescent-based formulation, the likelihood of an ARG is computed backward in time (from tips to the root), where a coalescent event collapses two existing lineages into one and a reassortment event ‘creates’ a new lineage from an existing one. This approach is analogous to the multi-species coalescent models used for joint gene tree–species tree inference [31, 32]. However, both approaches share the core assumption that the best path to the inference of reassortment events is via an ARG-like object, itself derived from incongruities between the segment trees.

**Figure 1:**
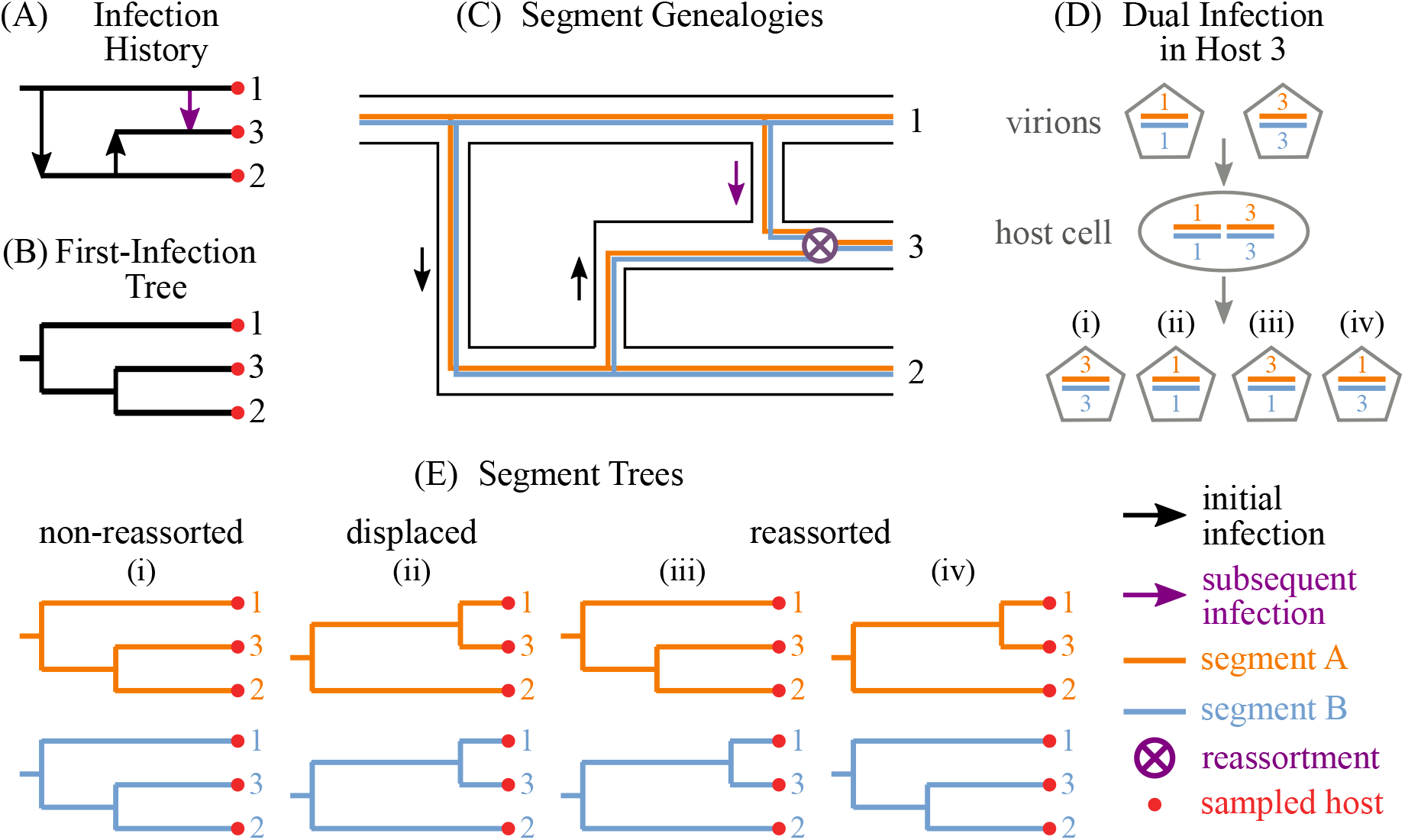
Dual infection can lead to reassorted segment trees. Time flows from left to right in panels A, B, C, and E. (A) The *infection history* encompasses all initial and subsequent infection events. In this small example, Host 1 infects Host 2, who then infects Host 3, and later Host 1 infects Host 3. Host 3 is then dually infected. (B) The *first-infection tree* includes initial infections of each host, but not subsequent infections or the identities of who infected whom. (C) The genealogy of each of the two viral genomic segments can be traced within the infection history. (D) Two virions in Host 3, the resident virion in Host 3 originated from the initial infection from Host 2 and the invasive virion from the subsequent infection from Host 1, can reassort upon dual infection of a cell within Host 3. (E) Depending on which virion is sampled from Host 3, the *segment trees* may show discordance with one another and/or with the first-infection tree. In outcome (i) the host maintains the viral genotype from its first infection, in (ii) the genotype from the second infection displaces the first, and in (iii) and (iv) a new reassorted viral genotype is generated.

In this paper, we argue that reassortment inference faces a fundamental problem: the number and pattern of reassortment events is not identifiable without reference to some aspects of the underlying, forward-time, epidemiological process, namely, a latent data structure that we call the ‘first-infection tree.’ We further identify four situations that we argue need to be explicitly handled by methods that infer reassortment. For this, we use a simple linear birth-death Markov chain simulation [33, 34] to show that these situations are common even in small samples with perfect data for very simple epidemiological models. By simulating sequence data from these models, we also assess the error properties of commonly used methods, showing that estimation of the number of reassortment events necessary to explain incongruities in the reconstructed segment trees is not consistently correct. Our approach to both identifying important problems and designing a simulation model was to simplify biological reality as much as possible to obtain a core set of problems that are not dependent on particular model formulations or biological details. We have also attempted to design our forward simulation models to be as consistent as possible with existing methods using identical model forms and parameterizations where possible. This work leads us to a few general conclusions: (1) the inference of viral genomic reassortment is not separable from the inference of epidemiological dynamics, (2) the ARG structure needs to be expanded to work for inference of viral genomic reassortment, and (3) coalescent models are a difficult starting point for inferring reassortment. Our objective is not to criticize existing methods or suggest that previous work is wrong; in fact, we argue that there are several paths to robust methods that can be obtained from modifications of existing open-source approaches. Rather, our intention is to clarify the conceptual challenges in inferring reassortment for future implementations.

## Results

Our results are laid out as follows. The first two sections illustrate logical problems that arise in the assessment of viral genomic reassortment. In those sections, we assume that all genealogies are binary and that reassortment is an instantaneous process that replaces one or more genomic segments in a given host with a copy from another extant host. In the final section, we use a forward linear birth-death simulation model to compute the frequency at which those problems arise in a theoretical study, and to simulate sequence and genealogical data to test methods for inferring viral reassortment. Details of the simulator and technical methods can be found in the Methods section and Supplemental Materials.

For clarity, we define a few standard and non-standard terms that we will use throughout the paper. We use the term *population history* to refer to the full history of all events that update the host dynamics or the genealogical structures in a population. In the context of a model, this is a record of who infected whom, when, and the times of all other sampling, removal, and reassortment events. Figure 1 illustrates several important subsets of the population history. The *infection history* shows which individual infected (or reinfected) which other individual, and at what time. The *segment trees* are the observed genealogies of the individual viral segments in a host population. The *first-infection tree* is the subset of the infection history that only includes the first infection event (*i*.*e*., the creation of a newly infected individual) for each infected and sampled individual, without the specific information of who infected whom at the internal nodes. The first-infection tree can be thought of as the counterfactual viral genealogy in the case where no reassortment occurred, in which case the first-infection tree and segment trees are all identical, regarding topology and branch lengths. An infection history defines exactly one first-infection tree, but one first-infection tree is consistent with many possible infection histories. We assume that the true genealogy is directly observable, though in reality this involves inferring time-scaled phylogenetic segment trees. Also, we make no attempt to deal with the intricacies of tree reconstruction since the difficulties with incongruence-based reassortment inference can be made evident without doing so.

### The fundamental problem

In this section, we illustrate the fundamental problem of reassortment using the hypothetical case where we have observed two segment trees and want to infer the true number of reassortment events required to explain their observed differences (Fig. 2(A)). Note that we make a distinction here between *visible* reassortment events that had some influence on the observed segment trees, and *invisible* reassortment events that are either not ancestral to the sampled viruses, or are ancestral but were replaced by a reassortment event that happened later in the same lineage (Fig. 2(B)).

**Figure 2:**
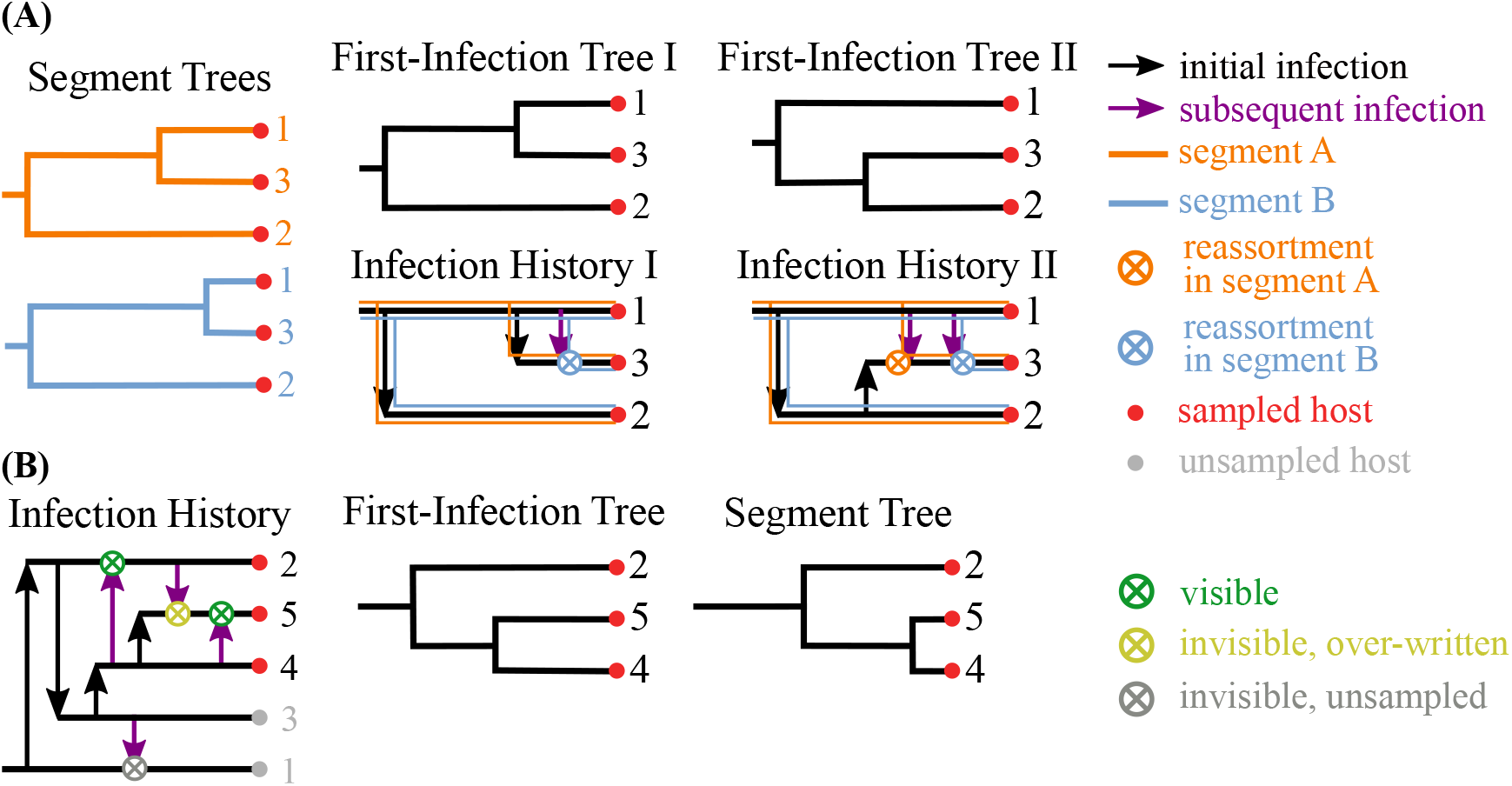
How the infection history, first-infection tree, and sampling process impact genomic reassortment inference. (A) Suppose that two segment trees are sampled. In a special case, First-Infection Tree I arose from Infection History I and is identical to one of the segment trees, so a single reassortment event is visible in the data. However, it could also be the case that First-Infection Tree II resulted from Infection History II, where two reassortment events are visible. Put another way, only one reassortment in segment B is needed to explain the segment trees’ difference from First-Infection Tree I, while one reassortment in each segment is need to explain why they both differ from First-Infection Tree II. (B) Of all the reassortment events in the full population (all four ⨂ shown), only some leave a visible imprint in the data. Reassortment events that occur in unsampled lineages are invisible, as are those that occur in sampled lineages but are over-written by another reassortment event later in the same lineage.

A common way of counting the number of visible reassortment events is to find the minimum number of ‘remove-and-rejoin’ operations [21, 26, 35] that are needed to reconcile the segment trees. The idea is to resolve any structural differences between segment trees by removing a tip or clade in one tree, along with its ancestral branch, and then rejoining that branch somewhere else in the same tree so as to reduce the differences between both trees. Figure S3 illustrates this idea. Generally, for such an approach the governing principle is parsimony: to propose the minimum number of reassortments that are needed to transform one segment tree into the other. Note that when we talk about incongruities in segment trees, we are referring to not only the tree topologies but also the branch lengths, which makes sense under the assumption that the segment trees can be observed without error. In practice, counting the number of reassortments might only rely on the inferred phylogenetic topology, under the assumption that differences in branch length are stochastic and/or due to evolutionary rate heterogeneity across tree branches. In that case, the remove-and-rejoin method is identical to the subtree-prune-regraft (SPR) method [35]. However, because our goal is to identify the fundamental problem to inferring viral genomic reassortment, we assume that reassortment events that change the branch lengths while preserving the topology in the segment trees are visible.

The problem that arises in applying remove-and-rejoin moves on the segment trees is that, even when applied correctly, it does not necessarily find all the reassortment events that are visible in the data when one also considers the underlying population history. Figure 2(A) illustrates the most basic case where the exact same set of segment trees can have a different number of visible reassortment events, depending on the infection history. While it is possible for a single remove-and-rejoin move (in this example) to convert one segment tree into the other, this resolution is not biologically possible under some infection histories. From this simple illustration, we obtain two important results: (1) reassortment events are not necessarily identifiable from segment trees without explicit reference to the unobserved population history, and therefore, (2) inference of viral reassortment is fundamentally linked to inference of epidemiological dynamics. That is, to know even the visible number of reassortment events in the smallest possible tree, we need some way of accounting for the probabilities of all possible infection histories (*e*.*g*., an epidemiological model).

We argue that finding the number of visible reassortment events in the observed data does not actually require knowledge of the full infection history; rather, the first-infection tree alone is sufficient. The first-infection tree can be thought of as the tree structure that would have been observed in the segment trees if there had been no reassortment. It therefore represents a starting point for calculating the number of reassortment events required to obtain each of the segment trees (Fig. S3). From this perspective, the minimum number of remove-and-rejoin moves is simply the minimum number of moves required to get from the first-infection tree to each of segment trees. Starting from a different first-infection tree can give a different minimum number of reassortment events required to explain the data. Without specifying a first-infection tree as a starting point, remove-and-rejoin methods under parsimony implicitly assume that one of the segment trees has the same structure as the first-infection tree. For example, in Fig. 2(A), unconditional remove-and-rejoin parsimony will find that only one move is necessary, but this is correct only under the assumption that the first-infection tree (I) has the same structure as segment tree A. In general, failing to specify a first-infection tree will bias results to fewer reassortments because it makes the assumption that one of the segments has no reassortment. Therefore, unconditional parsimony is inconsistent as a criterion for reconstructing reassortment. Without either knowing (or assuming) the first-infection tree or assuming some model that gives an uncertainty expressed as a probability mass over the space of first-infection trees, parsimony cannot be a valid criterion for inferring reassortment. The broader issue also extents to methods that use softer constraints such as a penalty based on the norm of the reassortment rates or a prior in the context of a Bayesian analysis without explicitly conditioning on or sampling first-infection trees.

### Four situations that make inference of reassortment dynamics difficult

In this section we illustrate four situations that cause further problems for estimating other quantities such as the reassortment rate and the time of reassortment events. We refer to these specific situations as ‘invisible’, ‘inaccurate’, ‘reversed’, and ‘obfuscated’, identifying where the potential for error arises in each case and the impact that it might have on reassortment inference. Any method that claims to be able to infer the history of reassortment in a population should be able to explicitly avoid or control for these errors.

Figure 3 illustrates the situation of *invisible* reassortment. This is the most straight-forward and easiest approach to correct situation. Invisible reassortment occurs when either a reassortment occurs in the part of the population that is unsampled, or when it is overwritten by another reassortment event that happens later (see Fig. 2(B)). In the context of a structural analysis of reassortment, this is not viewed as problem-atic inasmuch as such analyses attempt to identify the minimum number of reassortments which explain the data and would not claim to detect invisible reassortments. However, for methods that attempt to infer the reassortment rate, failure to correct this situation will bias estimates of the reassortment rate. In Fig. 3 the invisible reassortment shown is not ancestral to the sample and, therefore, will only directly influence inferences concerning the broader effect of reassortment in the population (*e*.*g*., the distribution of fitness effects caused by any reassortment). However, invisible reassortment also occurs along sampled lineages (*i*.*e*., ancestral to the sample). Along a given lineage and its descendants, we can only observe the most recent successful reassortment event(s). Figure S2 illustrates how we dismiss those invisible reassortments along the lineages before performing the remove-and-rejoin moves to resolve the structural differences between trees. If a method does not correctly integrate the probability that an inferred ancestral reassortment event is visible, it could lead to an underestimation of the reassortment rate or spurious support for heterogeneous reassortment rates. This is a particularly acute problem for methods based on the coalescent, as they condition on the observed data and are unable to resolve this problem as the sampling rate in such models is not defined.

**Figure 3:**
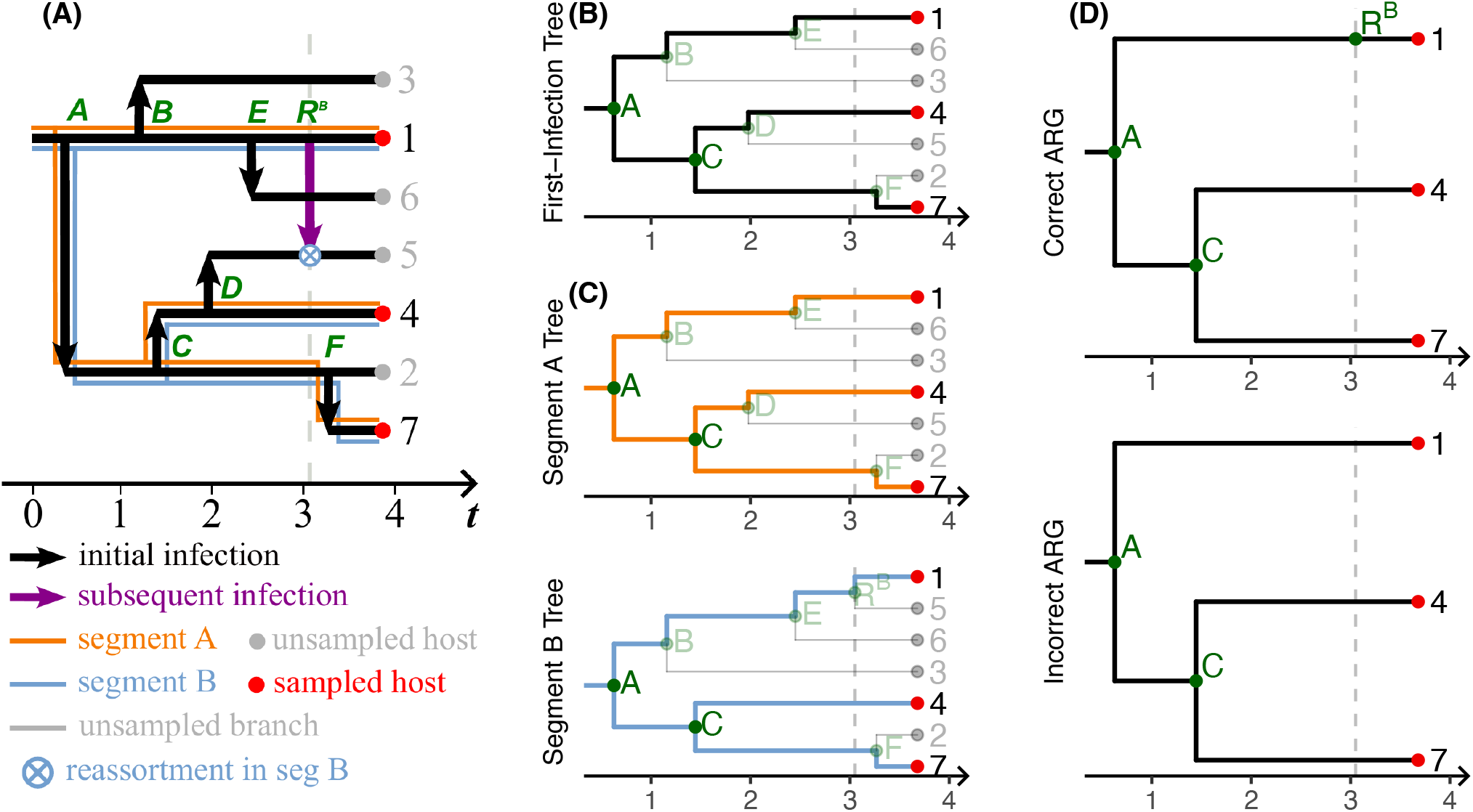
Invisible reassortment. This figure is an example of an *invisible* type of reassortment event that arises when a reassortment event either occurs in an unsampled part of the population or is replaced by a later reassortment event in the sampled part of the population. Trees are measured in arbitrary time units increasing from left to right. The full population history (A), first-infection tree (B), segment trees (C), and ARGs (D) are shown. The vertical grey dashed line indicates the time of that reassortment event.

Figure 4 illustrates the situation of *inaccurate* inference of reassortment times that occurs when both individuals (or one of their direct descendants) involved in a reassortment are sampled. At time *t*_*B*_, there is a reassortment event between samples 1 and 4, and, because both individuals are eventually sampled, the node corresponding to that event shows up as the most recent common ancestor of individuals 1 and 4 in the segment B tree. These types of nodes pose a problem for methods based on the manipulation of an ARG in the context of a reverse-time coalescent process. The situation is that in the coalescent-with-reassortment model, the coalescent and reassortment processes must be assumed independent to ensure that the coalescent prior makes sense. However, as is clear in this example and from the representation of reassortment events in our model, an internal node in one or more of the sampled segment trees can be caused by a reassortment event rather than a first-infection event. In Fig. 4, the node *R*^*B*^ corresponds to a reassortment event that produced a node in segment tree B, where both children of that node (individuals 1 and 4) were sampled leading to the node associated with the reassortment event being in the segment B tree. To explain this, a coalescent-based approach needs to insert two nodes into the ARG, one corresponding to the reassortment event to split and another separate node for that additional lineage to coalesce. Because both nodes cannot co-occur, a bias is introduced. In principle, the distance between these two nodes can be made arbitrarily small such that they effectively occur at the same time. However, this is unlikely in practice unless the data are highly informative (*i*.*e*., very long, clean sequences). In the more common case where the data cannot entirely overcome the coalescent priors, the distance between the two nodes will be influenced by the coalescent model which, by definition, will have some non-zero mean, encouraging bias in the estimation of the reassortment and coalescent times.

**Figure 4:**
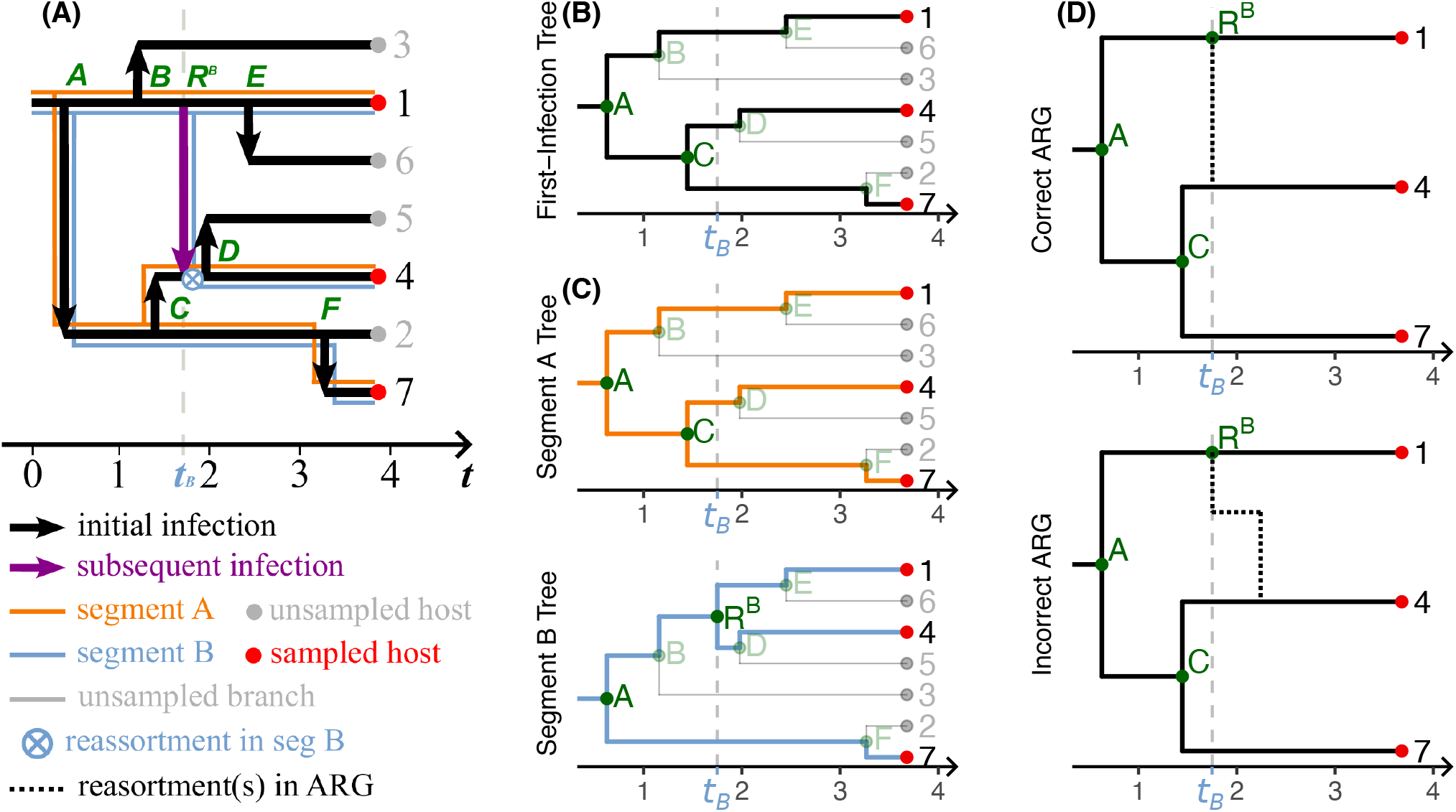
Inaccurate reassortment. This figure is an example of an *inaccurate* reassortment event that arises when both children of a reassortment event are sampled creating a node in the segment tree that was caused by reassortment and not a new infection. The full population history (A), first-infection tree (B), segment trees (C), and ARGs (D) are shown. The vertical grey dashed line indicates the time of that reassortment event and in (D), the black dotted lines indicate an actual or inferred reassortment in the population.

Figure 5 illustrates the situation of *reversion* in the inference of reassortment. Reversion occurs when a segment is passed between sampled individuals, but only one offspring of the reassortment event is sampled, leaving the node corresponding to the reassortment event to be unobserved in the segment tree (*i*.*e*., individuals 5 and 6 are unsampled in Fig. 5(C)). The statistical situation of reversion has a similar effect to that in inaccuracy, in that either one or both of the timing of the reassortment and corresponding coalescent events in an ARG will not be correct. The cause is slightly different, however. When only one lineage descending from a reassortment event is sampled, the corresponding ARG requires a tri-furcation (three lineages coming together in one node) at the sampled parent node of an unsampled reassortment event. Generally speaking, if some or all of the nodes of the ARG are assumed to follow a coalescent prior, then tri-furcations will not be allowed, creating a similar situation as in the case of inaccuracy, where additional, spurious nodes must be inserted into the ARG (nodes *A*^*′*^ and *C*^*′*^ in Fig. 5(D)).

**Figure 5:**
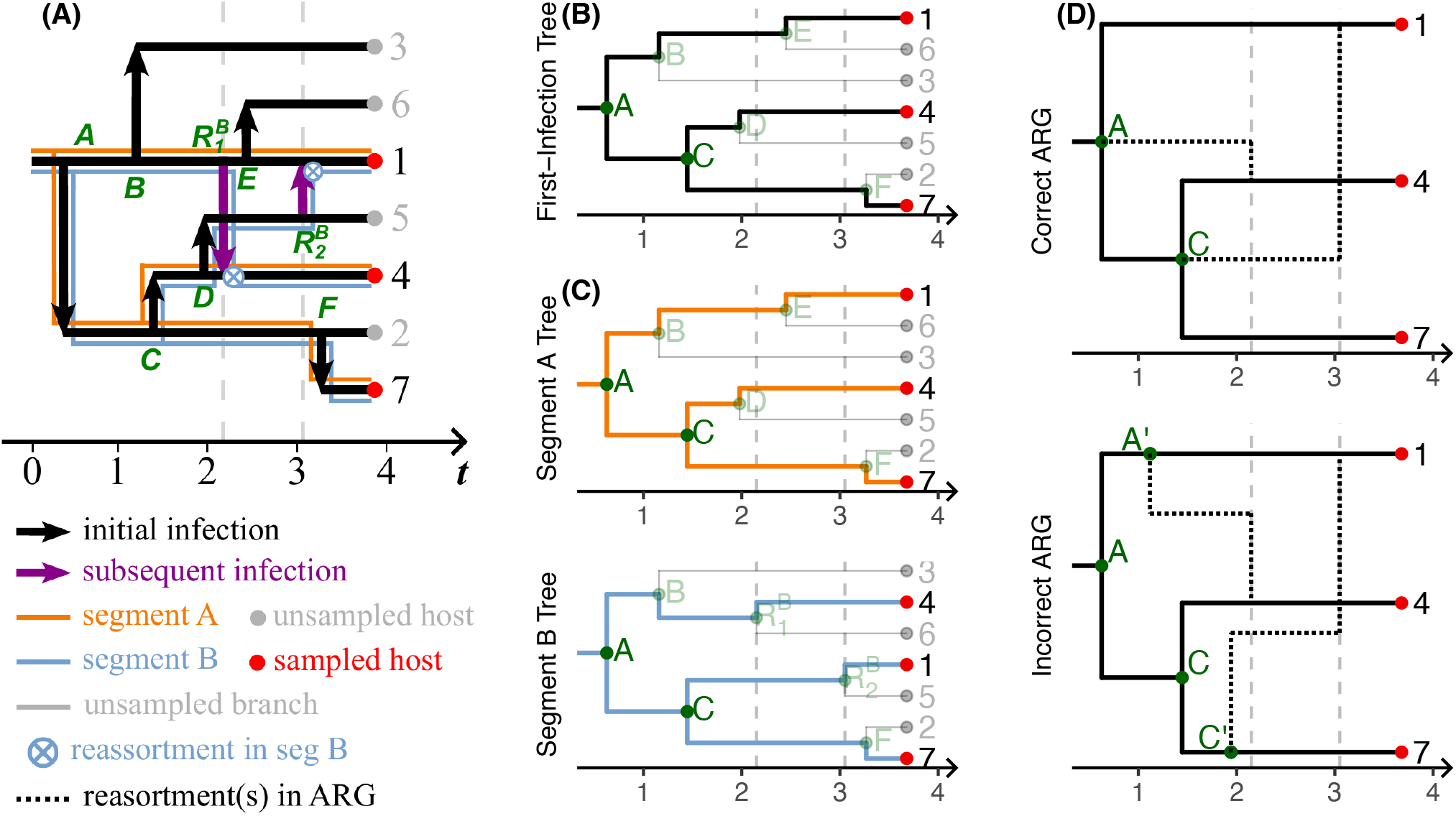
Reversed reassortment. This figure is an example of an *reversed* reassortment event that arises when only one child of a reassortment event is sampled creating the need for a trifrucating node in the ARG. The full population history (A), first-infection tree (B), segment trees (C), and ARGs (D) are shown. The vertical grey dashed line indicates the time of that reassortment event and in (D), the black dotted lines indicate an actual or inferred reassortment in the population.

Figure 6 illustrates the situation of *obfuscation* in the inference of reassortment. The defining aspect of obfuscation is that all segment trees have one or more reassortment events and at least one of these events is only partially sampled (*i*.*e*., only one of the descendants is sampled). We call this situation obfuscation because the first-infection tree no longer has the same structure as one of the segment trees. The effect of obfuscation is that the data appear to be explainable by fewer than the number of visible reassortments and the underlying population history is thus completely obscured.

**Figure 6:**
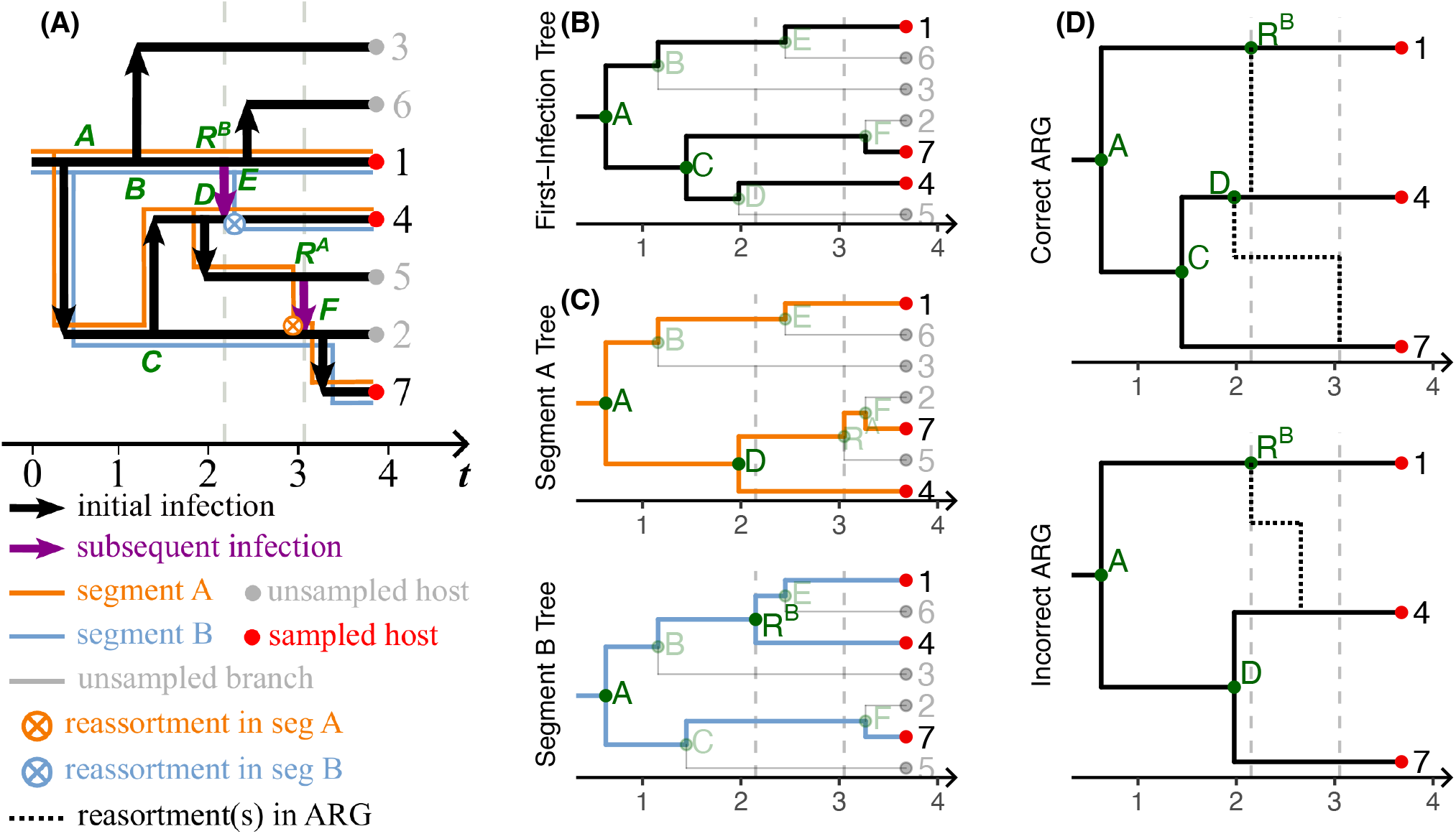
Obscured reassortments. This figure is an example of an *obscured* reassortment event that arises when both sampled segment trees contain reassortment events such that the first-infection tree no longer has the same structure as one of the segment trees. The full population history (A), first-infection tree (B), segment trees (C), and ARGs (D) are shown. The vertical grey dashed line indicates the time of that reassortment event and in (D), the black dotted lines indicate an actual or inferred reassortment in the population.

### The situations that cause problems for the inference of *visible* reassortment events are common

Of the four situations that we discussed above, three are potential problems for inferring visible reassortments. In this section, we assess the frequency at which those situations arise both theoretically and their impact on the accuracy of existing methods. We do not assess the frequency of invisible reassortment events as most reassortment events will be invisible, even in the context of an extremely large sample, and invisible reassortment events are not a problem in the assessment of visible reassortment events per se.

Using the population model detailed in the Methods section, we simulated a full epidemiological population history and extracted the sampled segment trees for a grid of different parameter combinations. We then counted the frequency of each of the different types of situations (details in the Supplementary Materials). Figure 7 shows the frequency of each situation for a range of simulation parameter values. In summary, the means of average frequencies of each situation are 26.1% (17%–45%), 6.7% (0.42%–9.6%), and 10.5% (0.49%–28.4%) for the situations *inaccuracy, reversion*, and *obfuscation*, respectively. The outcome variable is the proportion of nodes that have the type of referred situation averaged over 100 simulations. The *inaccuracy* situation increases as the first-infection rate becomes small with respect to a fixed sampling rate. This makes sense as inaccuracy occurs when both descendants from a reassortment event are sampled, which is more likely when the first-infection rate is low, meaning that a larger proportion of nodes in the tree are caused by reassortment events, rather than first-infection events. Both *reversion* and *obfuscation* follow a similar pattern of being less frequent when the first-infection rate is high and the reassortment rate is low with respect to the sampling rate, and more frequent when the first-infection rate is low and the reassortment rate is high. Both of these types of situations occur when the nodes caused by reassortment events are not fully sampled (*i*.*e*., they do not appear as a node in the sampled segment trees), which will occur with higher frequency when a larger fraction of the nodes in a tree are caused by reassortment events rather than first-infection events.

**Figure 7:**
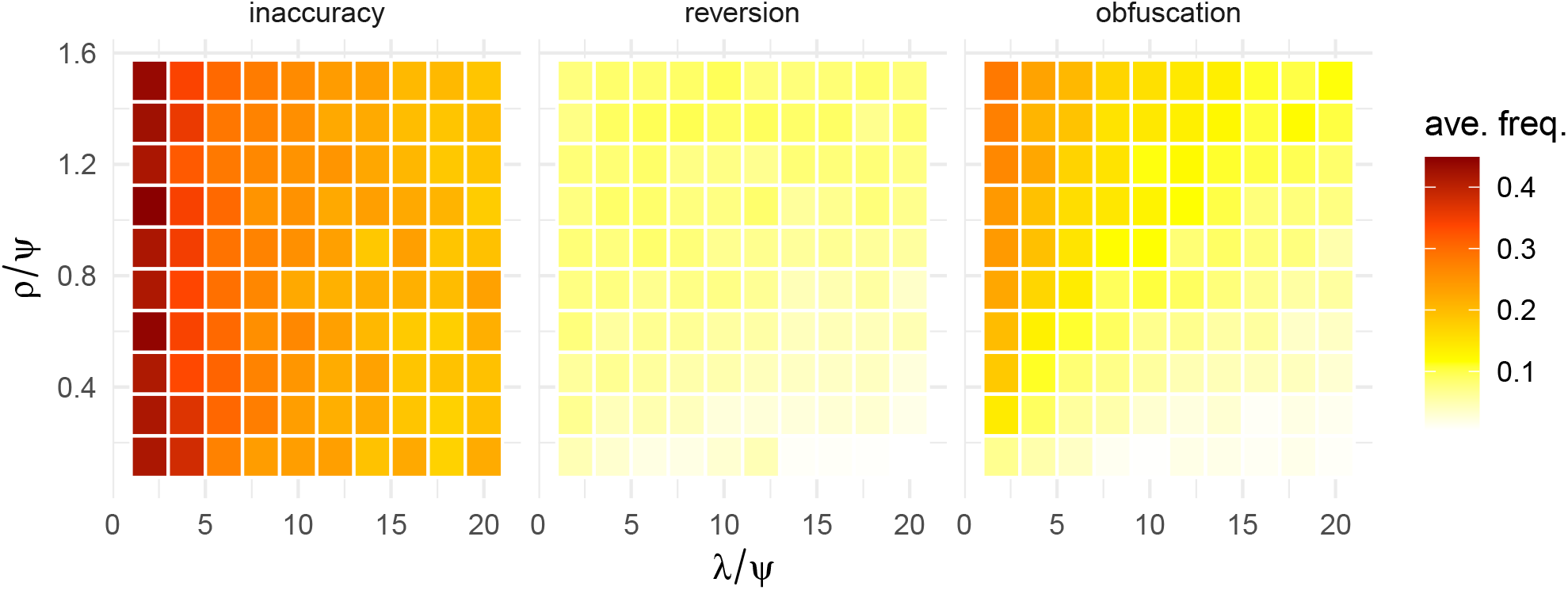
The frequency of the *inaccuracy, reversion* and the *obfuscation* situations. Starting from population size *I* = 100, the first infection rate is fixed at *λ* = 5 and the effective removal rate is *µ* + *Ψ* = 5, thus the expected population stays at *I* = 100. The total reassortment rate *ρ* is the sum of the rates for each segment and the joint reassortment rate: *ρ* = *ρ*_*A*_ + *ρ*_*B*_ + *ρ*_*AB*_, where *ρ*_*A*_ : *ρ*_*B*_ = 3 : 2 and *ρ*_*AB*_ = 0. Samples are taken sequentially at a constant rate *Ψ* until 50 individuals in the population are sampled.

Figure 8 shows the probability of inferring the number of visible reassortment events using two popular inference methods and an ‘omniscient’ approach that counts the observed minimum number of remove-and-rejoin moves in 100 simulated data sets for each condition. In this context, the remove-and-rejoin approach is called omniscient because we base it on the simulated population history rather than on inference, removing the issue of errors. In the case where reassortments only occur in one segment, the omniscient approach is always correct, by definition, but both the coalescent-based [27] and topology-based [22] approaches tend to get the right number of reassortment events about 15%–45% of the time. However, when both segments are allowed to reassort, the ‘omniscient’ method also tends to get the number of reassortment events wrong because one of the segment trees is no longer guaranteed to have the same structure as the first-infection tree. With the help of the simulated first-infection tree, the performances of both coalescent-based and topology-based methods are improved by about 10%. We further investigated the deviation in `CoalRe` [27] estimations of reassortment number and the reassortment timing across all the posterior ARGs in Figs. S4 and S5, respectively, showing that median errors in the assessment of reassortment times were generally about 12%–18% of the total tree height (ranging from about 5% to 60%). Adding the simulated first-infection tree reduces the median errors to approximately 10%, ranging from 4% to 65%.

**Figure 8:**
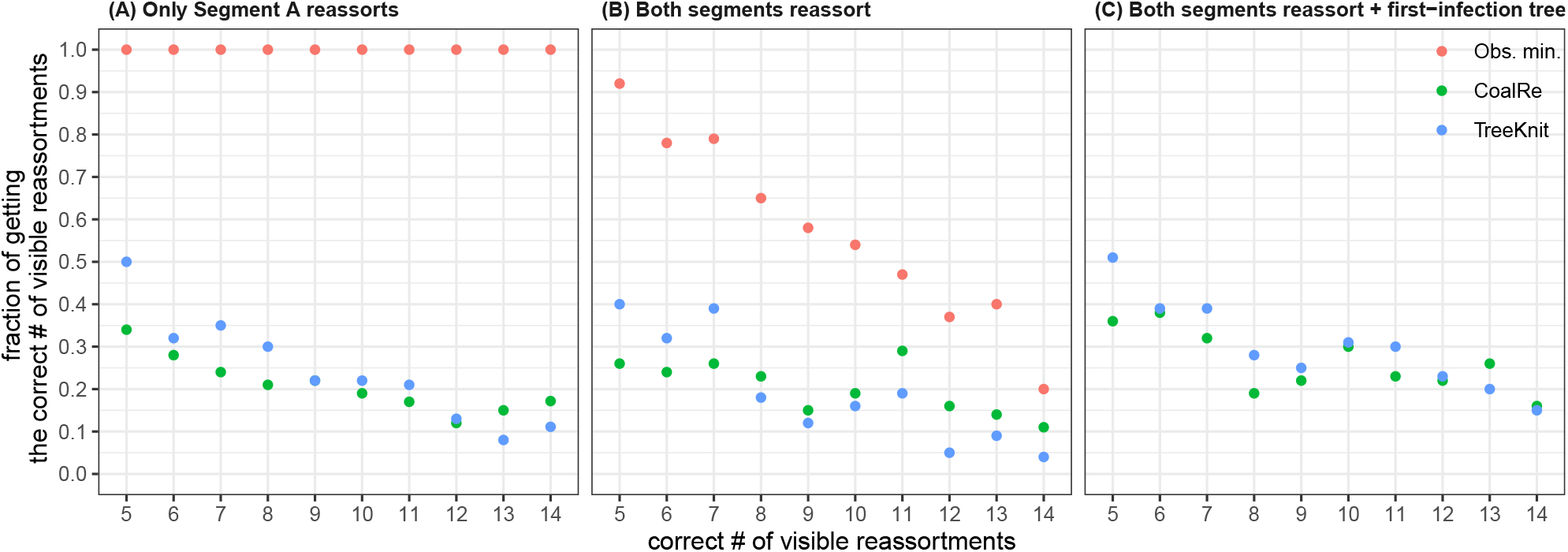
Fraction of correctly estimating the number of visible reassortments. With a constant expected population size *I* = 100 and rates *λ* = 5, *µ* = 4.75, *Ψ* = .25, and *ρ* = .2, we simulated 100 sets of trees with 50 samples for each of the number of visible reassortments, ranging from 5 to 14. For each set, we computed the observed minimum number of reassortments and estimated the number of reassortments by the inferred MCC ARG from `CoalRe` and the ARG from `TreeKnit`, such that we computed the fraction of 100 simulated trees that estimate the number of visible reassortments correctly. Two reassortment schemes are discussed: in (A), only segment A reassorts, *ρ*_*A*_ = .2 and *ρ*_*B*_ = *ρ*_*AB*_ = 0; in (B) and (C), both segments reassort but not simultaneously, *ρ*_*A*_ = .12, *ρ*_*B*_ = .08, and *ρ*_*AB*_ = 0. Two inference schemes are discussed: in (A) and (B), only segment trees are used for inference; in (C), segment trees and the simulated first-infection tree are used.

## Discussion

We attempted to detail what we believe to be the core issues in the inference of viral genomic reassortment and argued that these issues are common and cause problems for existing approaches. Given the importance of viruses with segmented genomes to global public health, we see resolution of these issues as being steps that lead to a more reliable understanding of the pandemic potential of those viruses.

Our analysis has important limitations to consider. Elements of our results, such as the sufficiency of the first-infection tree for resolving the inconsistency of parsimony, have not been proved or framed in a more rigorous context. While we believe these conclusions are likely to be proven true in the future, they should be understood to be contingent until more formal theories of the relationship between first-infection trees and the accuracy and precision of corresponding inferences can be established. Likewise, it is not so clear how the inconsistency of parsimony for the inference of viral genomic reassortment affects either penalized maximum likelihood methods or Bayesian methods that do not explicitly infer or condition on a first-infection tree.

An important high-level implication of this work is that the dynamics of infection and sampling in a population cannot be removed from the assessment of reassortment. That is, any claim about the history of reassortment implies a claim about the dynamics of infection that may be unreasonable or inconsistent with known facts. One of the ways that this makes reassortment a technically hard problem is that the coalescent framework that is popular in Bayesian phylogenetics [36] and phylodynamics [37] is not an ideal starting point for inferring reassortment. First, the sampling rate in the coalescent is not defined as a population parameter. The coalescent conditions on the sample and works backward in time, such that the sampling rate (or probability in a batch sample) is not meaningfully defined. Even if the sampling rate can be inferred post-hoc (*e*.*g*., by computing the implicit rate after-the-fact) given the inferred population size dynamics [38], it would still be insufficient for adjusting for unsampled reassortment events, because the coalescent is, by definition, silent on the population history where unsampled reassortment events would most likely reside. Second, the coalescent has no obvious way of incorporating reassortment events in the context of the coalescent process itself. The coalescent has been proved to be an efficient approximation for inferring from a single genealogy, in terms of forward-time formulations such as the Wright-Fisher, Moran, and linear birth-death models [29, 30, 39]. Reassortment is a coupled death-birth event that generates a new branch in a segment tree without changing the population size, and there is no theoretical foundation to link this forward-time model to the coalescent. Part of the reason that the coalescent is a popular formalism for Bayesian time-scaled phylogenetics is that it provides a theoretical basis for making, otherwise weakly identifiable, evolutionary rate models work. However, in the context of reassortment, some or even most nodes might have nothing to do with the population process assumed by the coalescent, and, therefore, are either wrongly included in the coalescent prior or are left with an implicitly ill-defined prior. Further, the coalescent with reassortment will, by necessity, not allow tri-furcations or a single node to be created by both a reassortment and coalescent event, which prevents such a method from addressing two of the four situations that we raised. Third, reassortment in a coalescent framework only makes sense when paired with a corresponding coalescent event (in reverse time, a reassortment node creates a new lineage in an ARG that will eventually need to coalesce), so both the reassortment rate and the coalescent process itself become uninterpretable. This is because the coalescent rate is, in most applications, computed as a function of the number of extant lineages and the effective population size. However, both first-infection and reassortment can result in a branching node in a tree, in which the former indeed increases population size, but the latter, formed by a coupled death-birth of an individual, does not change the population size. This convolution of the coalescent event and the reassortment event confounds the interpretation of their respective rates.

We argued that a reliable method for inferring reassortment both in the number and timing of events needs to address the fundamental situation with under-counting and the specific statistical situations that we raised. To our knowledge, no existing software or proposed method meets these criteria, but there is very well-developed software that solves many of the most difficult aspects of the problem already. We see three practical ways that already-published methods could be adapted to address some or all of these situations.

We demonstrated that finding the maximum parsimony number of remove-and-rejoin moves between sampled segment trees is not a consistent way to count the correct number of visible reassortment events, unless the first-infection tree is known. Existing topology-based methods (like those implemented in `TreeKnit` [22]) could be adapted to search the space of remove-and-rejoin moves for a set of segment trees given a first-infection tree. It is unlikely that the first-infection tree would be known, so a more reasonable way to deal with conditioning on the first-infection tree is to treat it as a nuisance variable that we would need to integrate out in the context of some prior distribution over the space of first-infection trees. Simulation methods could be used to generate first-infection trees from some model parameters, and existing remove-and-rejoin algorithms could be applied to all pairs of first-infection tree and segment tree. Effectively this would both correct for the under-counting issue and would allow integration of prior knowledge in terms of the simulation model assumptions and the prior on the model parameters. Realistically, however, this approach might require a more theoretically robust understanding of how epidemiological dynamics constrain the inference of reassortment to make the computations feasible.

Methods that jointly search both the tree space and model parameter space provide not only a more complete reporting of uncertainty, but also direct estimates of the timing of reassortment events. The BEAST2 package `CoalRe` already implements most of the necessary functions that would need to be used to modify the process of sampling the joint posterior of first-infection trees and segment trees rather than an ARG-like object in the context of a prior based on a forward-time model of epidemiological dynamics and sampling. Further, this approach could be adapted to model non-linear epidemiological dynamics. We discuss the mathematical details of how this might work in the Supplemental Materials.

This paper’s point could be summarized as, “to understand reassortment, one must place it in the context of a full population process.” The problem is that there is no currently known method for computing the likelihood of either a sequence alignment or a set of segment trees given the population models that we talk about in this paper. The issue lies in proposing the candidates of latent sampled first-infection trees and computing their likelihoods, which may be computationally challenging. Therefore, one potential way to deal with this is to treat the mapping between a population model and observable data through the lens of likelihood-free methods, such as Approximate Bayesian Computation (ABC) [40]. These methods work by defining a metric space based on some set of statistics computed on the observed data and data simulated from the population process of interest. Recently Voznica *et al*. [41] proposed a deep-learning approach along these lines where, rather than picking a set of statistics by hand, they allow a neural network to approximate a mapping between model parameters and simulated tree structures for determining the population model structure, estimating the parameters, and exploring the uncertainties. A similar approach might prove fruitful for fitting complex models with genomic rearrangements such as reassortment.

A final consideration for the use of population models, as both explicit theories and data analytic tools, is that a well-defined population model can serve as a common domain for scientific hypotheses and the statistical tests needed to probe those hypotheses. Consider that genomic reassortment can occasionally produce highly-fit viral strains that then rapidly spread in a population. It is easy to imagine how one might integrate this question into a population model, *e*.*g*., having a neutral type and an advantageous type (in terms of increased transmission rate) of reassortment. If we were able to fit that model to data, we could characterize a direct test of our scientific question as a clearly defined statistical test of the model (*e*.*g*., whether or not one transmission rate is larger than the other).

## Methods

This section provides detailed interpretations and justifications on defining reassortment as a coupled death-birth event, incorporating reassortment in a simple population model with given parameterization, and simulating trees and sequences.

### A conceptual operationalization of reassortment in the context of a population process

Reassortment is a complex process that involves cellular co-infection, competition among different viral strains, host immune responses, and the biophysics of genome packaging and virion formation. Each host-virus system will be unique in ways that may affect inferences regarding, for example, the timing of reassortment events. However, taking out the complex biology and within-host dynamics, the most basic operationalization of reassortment in a model is that of a coupled death-birth event. We illustrate how reassortment changes the evolutionary histories of viral segments in Fig. S1 for a hypothetical virus with two segments, A and B. In that case, host 0 co-infects host 2, swapping the B segment. Host 2’s original B segment is lost in the population and their original branch in segment B is dissolved (*i*.*e*., the original branch ‘dies’). Correspondingly, host 2 now has a segment B that is directly descended from host 0 (*i*.*e*., a ‘birth’ occurs in segment B at host 0). This type of coupled death-birth event is how reassortment produces conflicts in both topology and branch lengths in segment trees, due to differences in their evolutionary histories [42]. Understanding the abstract effect of reassortment on genomic segment trees as this type of coupled death-birth event, in the context of a broader population process, is central to understanding the situations we raise in this paper.

This model defines a population process that consists of both the epidemiology of the host population and the ancestry of the viral population. Individuals can become infected and, once infected, can become infected again. We make the simplifying assumption that subsequent infection is resolved instantly, such that an individual only hosts one viral genotype at a time. We reserve the term *re-infection* for the situation in which the new viral genotype within the host differs from the previous one. To keep the model simple, we define reassortment as the combination of re-infection, cellular co-location, reassortment of one or more genomic segments, and dominance of the new reassortant at the host level. Each subset of genomic segments can, in principle, have a unique reassortment rate due to differences in within- or between-host survival, or the biophysical aspects of viral particle formation. Infected individuals leave the system by either dying or becoming sampled. For simplicity, we assume sampling is destructive to avoid the important but out-of-scope issues that arise in sampling direct descendants of already-sampled tips in phylogenetic trees. We assume all individuals are exchangeable, so there is no host- or virus-level heterogeneity, such as different behaviors or fitness.

### Simulating the frequency of inaccuracy, reversion, and obfuscation

We implemented the conceptual model as a linear birth-death Markov process [33, 43] using the suggested formalism for simulating genealogical processes in [39]. The model contains two distinct levels of biological processes: epidemiology of the host population and ancestry within the viral population (Table S1). A conceptual description is provided here, and a more complete technical description is available in the Supplementary Materials and Ref. [39]. The model assumes a viral infection with two segments (labeled A and B) and has six per-capita rates: the first-infection rate, *λ* ; the recovery rate, *µ*; the sampling rate, *Ψ*; the reassortment rates for segment A and B, *ρ*_*A*_ and *ρ*_*B*_, respectively; and the joint reassortment rate, *ρ*_*AB*_, accounting for the simultaneous reassortment or the displacement as shown in Fig. 1(E) option (ii). Note that the effective removal rate of an individual is *µ* + *Ψ*, as we assume infected individuals who got sampled are immediately hospitalized or quarantined and thus removed from the infected population. The spread of infection in the host population is described by stochastic exponential growth at rate *λ* − (*µ* + *Ψ*). The model records the full infection history (identity of source and recipient, and time of infection) in a format that enables much of the downstream computations that we use to count types of reassortment inference situations. The first-infection tree is a subset of infection history that includes only the sub-tree of sampled tips and the nodes corresponding to the time they were first infected; source and recipient identity are also removed.

The descriptions above apply to the entire population of hosts and the viruses that infect them, *i*.*e*., the full population process. Not all individuals will be observed, however, and the sampled genealogy only represents the ancestral relationships between viral genomic samples. In the simulation, the full infection history, first-infection tree, and segment trees are pruned to include only the samples taken. Samples are taken sequentially from the infected population at the per-lineage sampling rate, *Ψ*. For visual simplicity, figures illustrating stylized population histories or trees are always shown as if a homochronous, batch sample was taken at a given time.

All simulations start from one single infected individual (*I* = 1), while setting the first-infection rate *λ* = 200 and shutting down recovery, sampling, and reassortment until the population size reaches *I* = 100. We then turn on reassortment (*i*.*e*., *ρ*_*A*_ > 0 or/and *ρ*_*B*_ > 0, *ρ*_*AB*_ = 0) and run a burn-in period where the birth and removal rates are equal but there is no sampling (*i*.*e*., *λ* = *µ* = 5 and *Ψ* = 0) until the root time of the tree is later than the time of reaching *I* = 100, before proceeding with sampling (*i*.*e*., *λ* = *µ* + *Ψ*, where *µ* > 0 and *Ψ* > 0). We run the simulation until 50 samples are obtained. This approach guarantees that the expected population size stays constant at 100 during the studied period of the obtained trees. For each situation discussed in Results, we produced 100 stochastic replicates and computed the average frequency for each combination of parameters.

### Simulation and analysis of sequence data

We first simulated the full population history such that the expected population size is (*I* = 100), with rates *λ* = 5, *µ* = 4.75, *Ψ* = 0.25, *ρ* = *ρ*_*A*_ + *ρ*_*B*_ + *ρ*_*AB*_ = 0.2, until 50 samples were obtained. These parameters are chosen to align with simulations in previous literature [22, 27], meanwhile making sure the frequencies of aforementioned situations are significant enough. For each of the number of visible reassortments, ranging from 5 to 14, we produced 100 replicates. Each segment genealogy represents precisely the history of that segment, but inferring a segment tree in practice requires genetic sequence data. Therefore, we used IQ-TREE 2 [44] to simulate alignments with a sequence length of 10^3^ nucleotides using the Jukes-Cantor model for each tree, at an evolutionary rate 5 *×* 10^−3^ per site [27].

We analyzed our simulated data with two leading software packages for inferring reassortment. To use the `CoalRe` phylogenetic network model, we imported the alignments into BEAST2 and used the same mutation model. `TreeKnit` only requires the Newick format of tree data, so we directly input each set of trees.

Two different inference schemes were used: (1) using segment trees as input data only, and (2) using the simulated first-infection tree and the segment trees as input data. In `CoalRe`, the simulated alignments for the first-infection tree were with a length of 10^4^ nucleotides, approximately fixing its tree structure. In `TreeKnit`, we had pair-wise inputs, the first-infection tree and Segment A tree and the first-infection tree and Segment B tree, and summed up the number of reassortment events estimated separately from these inputs to have the total number of estimated reassortments.

## Acknowledgements

This work was supported by grants from the U.S. National Institutes of Health (Grants #R01AI167048 to CMP, #R01AI087520 to TL, #R01AI143852 to AAK), and by the Joint U.S. National Science Foundation/National Institutes of Health Interface Program (Grant #1761603 to AAK).

## Supplementary Materials

### Markov Genealogy Process with reassortments

Two essential elements for a Markov Genealogy Process with reassortment are: (a) a population process *𝒳*_*t*_ that describes the population dynamics, and (b) a segmented genealogy process 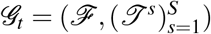 that incorporates the viral evolution along with the population dynamics, where *S* is the fixed number of viral genome segments. For a minimal model structure (*i*.*e*., a linear birth-death-sample model) for a two-segment virus, we can define: (a) the population process as *𝒳*_*t*_ = (*I, G, M*^*A*^, *M*^*A*^), where *I* is the infected population, *G* is the sample size, and *M*^*A*^ and *M*^*B*^ indicate the cumulative number of reassortments occurring in segment A and B, respectively; and (b) the genealogy process *𝒢*_*t*_ = (*ℱ, 𝒥* ^*A*^, *𝒥* ^*B*^), where *ℱ* denotes the first-infection tree, and *𝒥* ^*A*^ and *𝒥* ^*B*^ denote the segment A and B trees, respectively. Table S1 summarizes how the four most important events, *i*.*e*., first infection, recovery, sampling, and reassortment, change the dynamics of the population *𝒳*_*t*_ and the genealogies *𝒢*_*t*_. Theoretical details can be found in [1].

### Implementation details on dismissing invisible reassortments

In a simulated tree, an orange node marks a reassortment that has occurred in the other segment tree(s) and the first-infection tree, and a gold node denotes a reassortment that has occurred in the respective segment tree. Figure S2 left panels show a raw first-infection tree and the corresponding raw segment tree, before dismissing invisible reassortment nodes. The first-infection tree only contains orange reassortment nodes because no reassortment is allowed in the first-infection tree, while the segment tree only contains gold reassortment nodes because only this one segment reassorts in this example. We highlight the discording branching nodes in light green and purple shades, where purple shades denote the reversed branching nodes. Reassortments can become invisible in the sampled tree due to later reassortments along the same branch or on the same clade. For instance, in the first-infection tree at the left panel of Fig. S2, reassortment node No. 75 is invisible due to No. 117, and No. 15 is invisible due to No. 58 and No. 117. By dismissing those invisible reassortment nodes in both trees, we can have the minimum number of reassortments that correspond to the observed structural difference between the first-infection tree and the segment tree, as shown in the Fig. S2 right panels.

### The visible reassortment events and the remove-and-rejoin table

Figure S3 shows how one performs the ‘remove-and-rejoin’ method and counts the minimum number of reassortments given the first-infection tree and segment trees. After dismissing all invisible reassortment nodes, one can conduct the ‘remove-and-rejoin’ moves by comparing the first-infection tree and a segment tree. For example, dissolving the clade associated with reassortment node *a*_1_ removes branching node *D*, while rejoining the whole clade as one of the descendants of the node *B* creates a new branching node 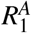, and eventually reduces the structural differences between the first-infection tree and the segment A tree. By repeating these remove-and-rejoin moves between the first-infection tree and each of the segment trees, one can write down two ‘remove-and-rejoin’ tables. Note that, node *c*_1_ implies that the corresponding reassortment occurs in both segments A and B simultaneously. This reassortment only causes a difference in branch length between the first-infection tree and the segment trees, while excluding which may result in biased estimation in mutation rates for reconstructed phylogenetics. Combining two tables and eliminating the duplicates gives us minimal remove-and-rejoin moves and the visible number of reassortments. With the help of the remove-rejoin table, one can produce the correct ARG.

### The number and frequency of three situations among visible reassortments

Using the simulated segment trees, one can easily identify the inaccuracy situation by counting the number of reassortment nodes that are aligned with its ancestral branching nodes (Table S1). For example, in either the segment A tree or segment B tree in Fig. S3, reassortment node *c*_1_ is aligned with the new branching node 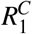, indicating the reassortment time of *c*_1_ is identical to the branching time of 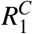. The reassortment node *c*_1_ falls into the *inaccuracy* situation category.

Reversion indicates that a seemingly consistent branching node in both trees, *i*.*e*., same node label and branching time, actually has distinguished descendants in two trees. For instance, branching node *D* appears in both the first-infection tree and the segment A tree in Fig. S3, with different descendant tips. We then call node *D* a reversed node, and the count for the *reversion* situation between the first-infection tree and the segment A tree is one.

Obfuscation is the most abstract situation among the four proposed, however, the easiest to identify. One can produce the visible remove-rejoin table by comparing the first-infection tree and each of the two segment trees, as shown in Fig. S3. The same procedure can be performed with only segment trees, assuming one of them is identical to the first-infection tree, such that we can have an “observed” remove-rejoin table. Extracting the true remove-rejoin table by the observed remove-rejoin table helps identify the degenerated moves.

The frequency of problematic situations can be computed by dividing the counts of the respective situation by the number of visible reassortments. For instance, in Fig. S3, the number of visible reassortments is 4, among which reassortment *c*_1_ corresponds to the *inaccuracy* situation, reassortment node *a*_1_ or *a*_2_ corresponds to the *reversion* situation. We thus have the frequency of the *inaccuracy* situation is 1/4 and that of the *reversion* is also 1/4.

### Adding the first-infection tree to the ARG likelihood function

For existing coalescent methods, such as `CoalRe`, to address the situations we identified in this paper, the first-infection tree would have to be incorporated in the likelihood function along with the segment trees. The classic Felsenstein likelihood [2] of the segment trees is then the last ingredient in the posterior distribution. We can then write

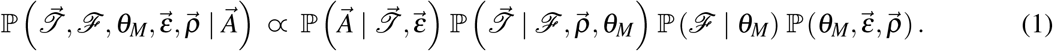

Here, *θ*_*M*_ is the set of parameters for a given population model *M*; *ℱ* is the first-infection tree induced by the given model *M* and parameters *θ*_*M*_; elements in vectors 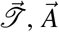, and 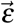 represent the corresponding segment tree, alignment, and mutation model, for the respective segment, separately; and 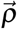 are the reassortment rates for each and all combinations of segments. The expression on the right-hand side consists of the likelihood of segment trees 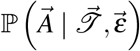, the segment trees prior distribution given remove-and-rejoin tables from the first-infection tree 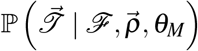, the first-infection tree prior distribution ℙ (*ℱ* | *θ*_*M*_), and the joint parameter prior distrubtion 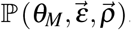. Specifically, by including both the topological difference and the difference in branch length between, the first-infection tree and the segment trees in the remove-and-rejoin tables, that is in 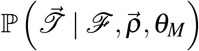, play a role in inferring the mutation model 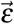.

**Table S1:**
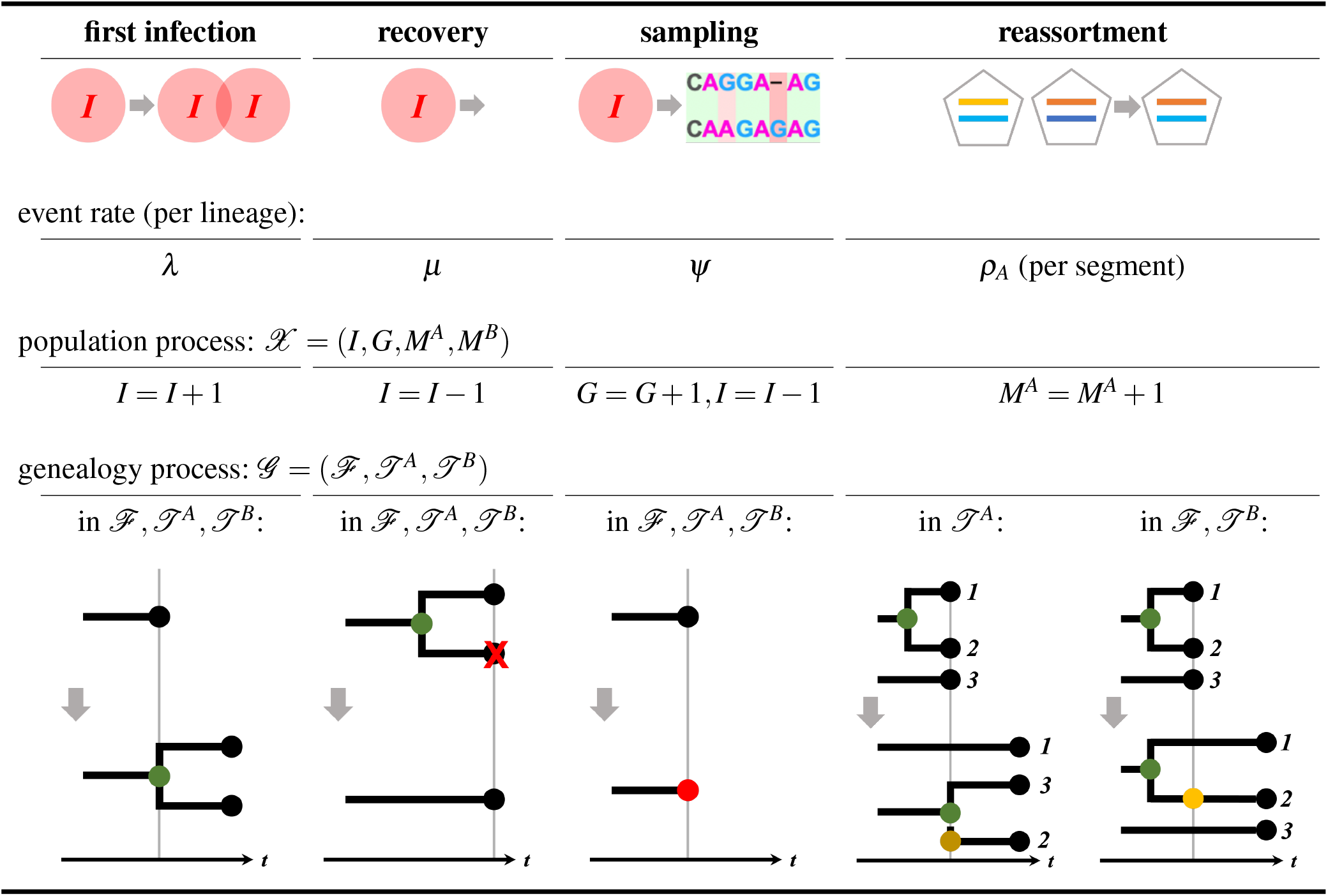
Summary of the population process, the genealogy process, and the important events. The population process *𝒳* is framed into a birth-death-sample model, represented by the number of infected population *I*, the cumulative count of samples *G*, and the cumulative counts (*i*.*e*., *M*^*A*^ and *M*^*B*^) of reassortments of Segment A and B, respectively. The genealogy process *𝒢* consists of three genealogies, *ℱ, 𝒥* ^*A*^, *𝒥* ^*B*^. Four important events and their impact on *𝒳* and *𝒢* are summarized: (a) a first infection event increases *I* by one and introduces a new branch and a new lineage in all three genealogies in *𝒢*, at rate *λ* ; (b) a recovery event reduces *I* by one and removes a tip and its corresponding branch from all genealogies, at rate *µ*; (c) a sampling event increases *G* but decreases *I*, and marks a tip as a sample then terminates it in all three genealogies, at rate *Ψ*; and (d) a reassortment event in segment A increases the count *M*^*A*^ by one, then performs a death-birth operation, *i*.*e*., dissolve the branch and re-create the lineage as a new branch, in only *𝒥* ^*A*^, marking the respective lineage by a gold node, while marking the same lineage in *ℱ* and *𝒥* ^*B*^ by an orange node to demonstrate the correspondence, at rate *ρ*_*A*_.

| first infection | recovery | sampling | reassortment |  |
| --- | --- | --- | --- | --- |
| event rate (per lineage): |  |  |  |  |
| $\lambda$ | $\mu$ | $\psi$ | $\rho_A$ (per segment) | |
| population process: $\mathcal{X} = (I, G, M^A, M^B)$ | | | | |
| $I = I + 1$ | $I = I - 1$ | $G = G + 1, I = I - 1$ | $M^A = M^A + 1$ | |
| genealogy process: $\mathcal{G} = (\mathcal{F}, \mathcal{T}^A, \mathcal{T}^B)$ | | | | |
| in $\mathcal{F}, \mathcal{T}^A, \mathcal{T}^B$: | in $\mathcal{F}, \mathcal{T}^A, \mathcal{T}^B$: | in $\mathcal{F}, \mathcal{T}^A, \mathcal{T}^B$: | in $\mathcal{T}^A$: | in $\mathcal{F}, \mathcal{T}^B$: |

**Figure S1:**
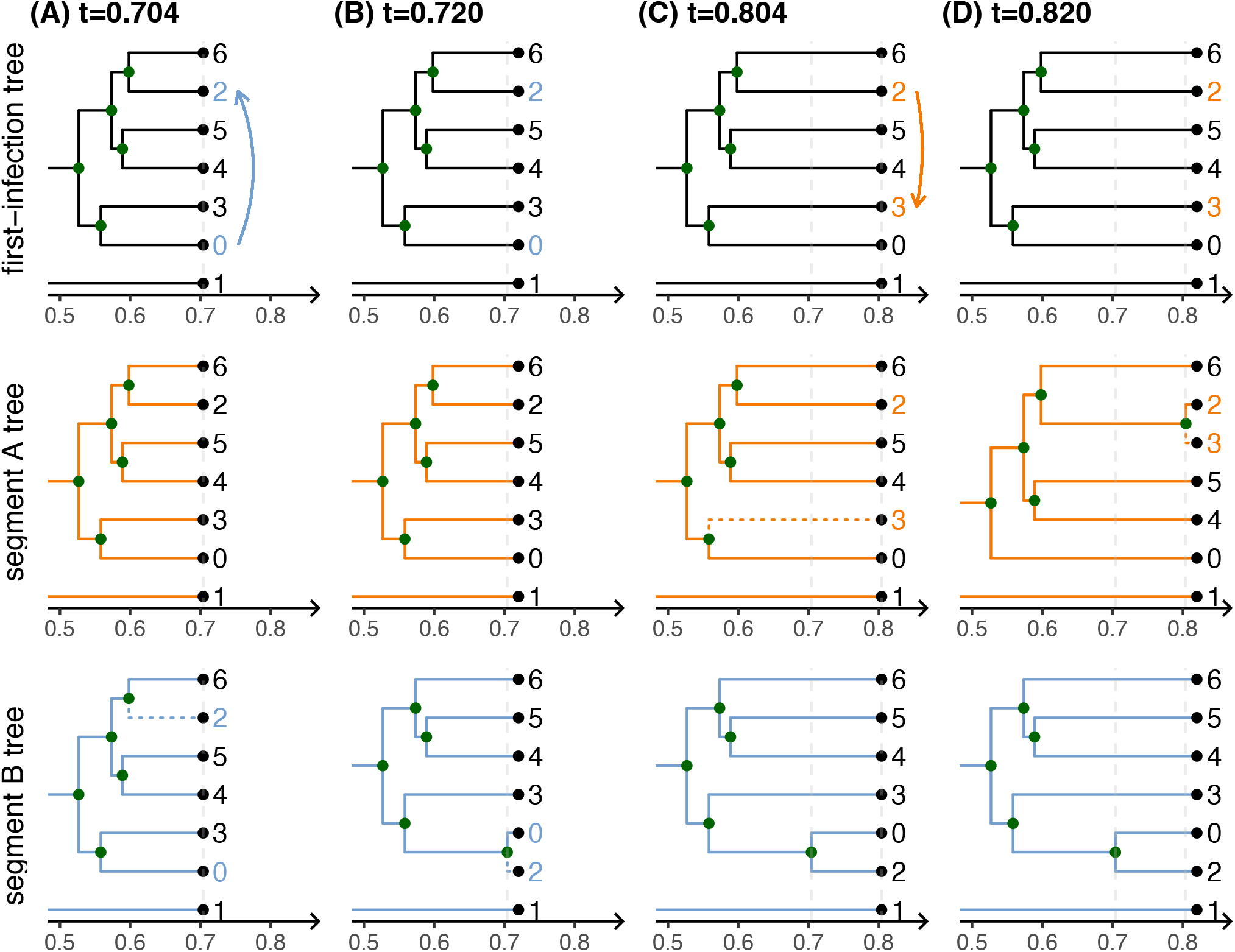
Reassortment events in the genealogy process for a two-segment virus. Two reassortment events occur during the demonstrated period, from *t* = 0.704 to *t* = 0.820. (A) At *t* = 0.704, the donor, host 0, re-infects the recipient, host 2, resulting in the swapping of genetic materials in segment B between hosts 0 and 2; (B) this first reassortment event in segment B will be reflected by the death of host 2 followed by the rebirth of host 2 in host 0 in segment B tree, and no other events occur till *t* = 0.720 and afterward; (C) at *t* = 0.804, host 2 re-infects host 3, resulting in swapping of genetic materials in segment A between hosts 2 and 3; (D) the second reassortment event in segment A will be reflected by the death of host 3 followed by the rebirth of host 3 in host 0 in segment A tree, and no other events occur till *t* = 0.820. Arrows indicate the re-infections, starting from the donor to the recipient. The blue arrow represents reassortment event induced by re-infection in segment B, and the orange one, in segment A. Dashed branches show the death-rebirth of the recipient in the corresponding segment tree, defined as a reassortment event.

**Figure S2:**
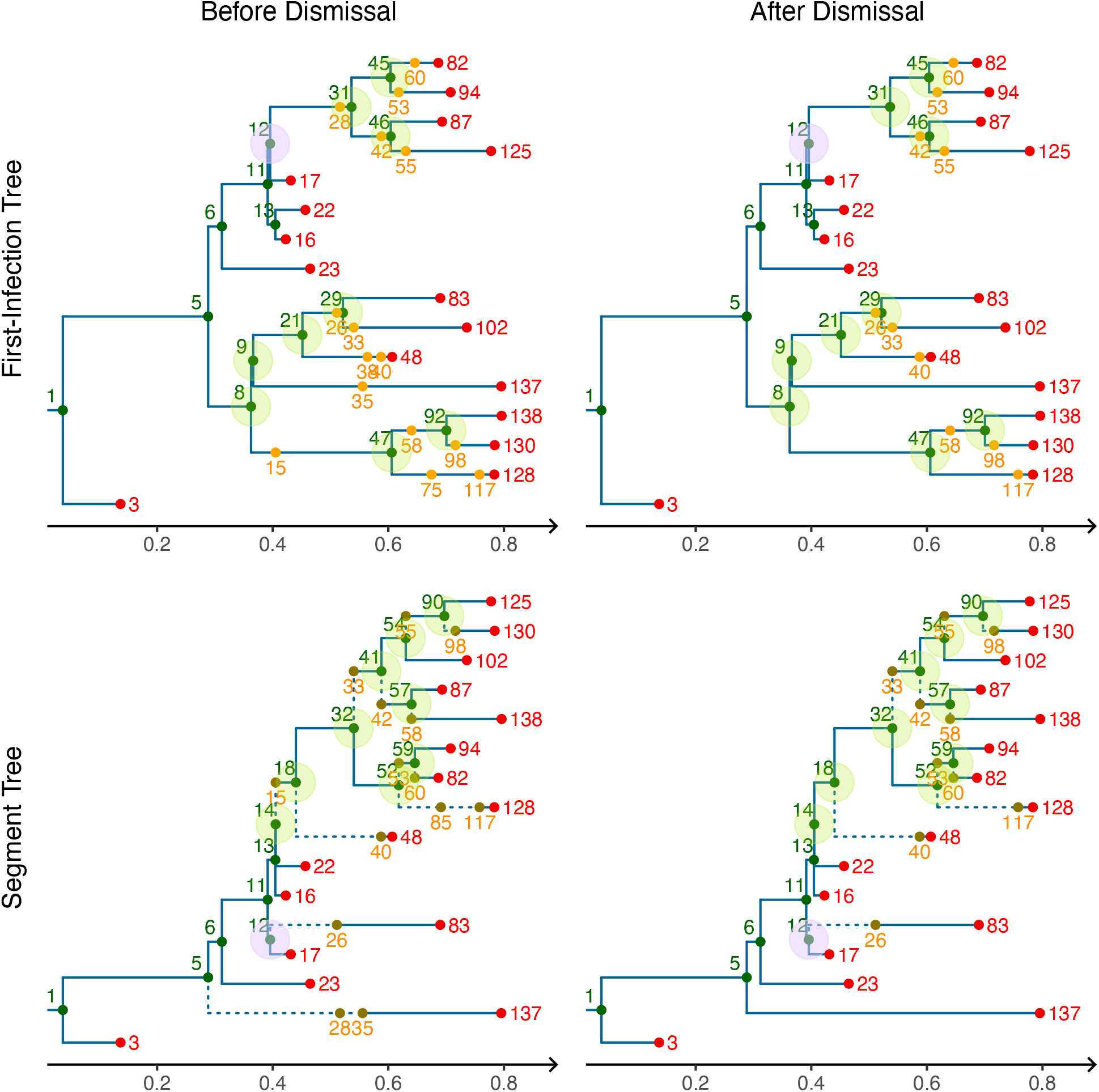
Dismiss invisible reassortments. This figure serves as a demonstrative example of the Dismissal Procedure. Our implementation records every node/tip and its label in the trees: shown in green, red, or orange/gold in the figure. Specifically, reassortment events are indicated by orange nodes in the first-infection trees and gold nodes in the segment tree that conducts the corresponding reassortments. Discording branching nodes are highlighted in green shape and the reversed branching nodes are in light purple shape. Before dismissal, in total 15 reassortment events are preserved in the sample trees, but some of them are not shared by both trees or in fact overwritten by the latter ones and will never be observed among samples. After dismissing these invisible reassortments, in total 10 reassortments were left, corresponding to 9 discording branching nodes and one reversed branching node between the first-infection tree and the segment tree. The remained reassortments are the minimum that is responsible for the discording and reversed branching numbers between trees.

**Figure S3:**
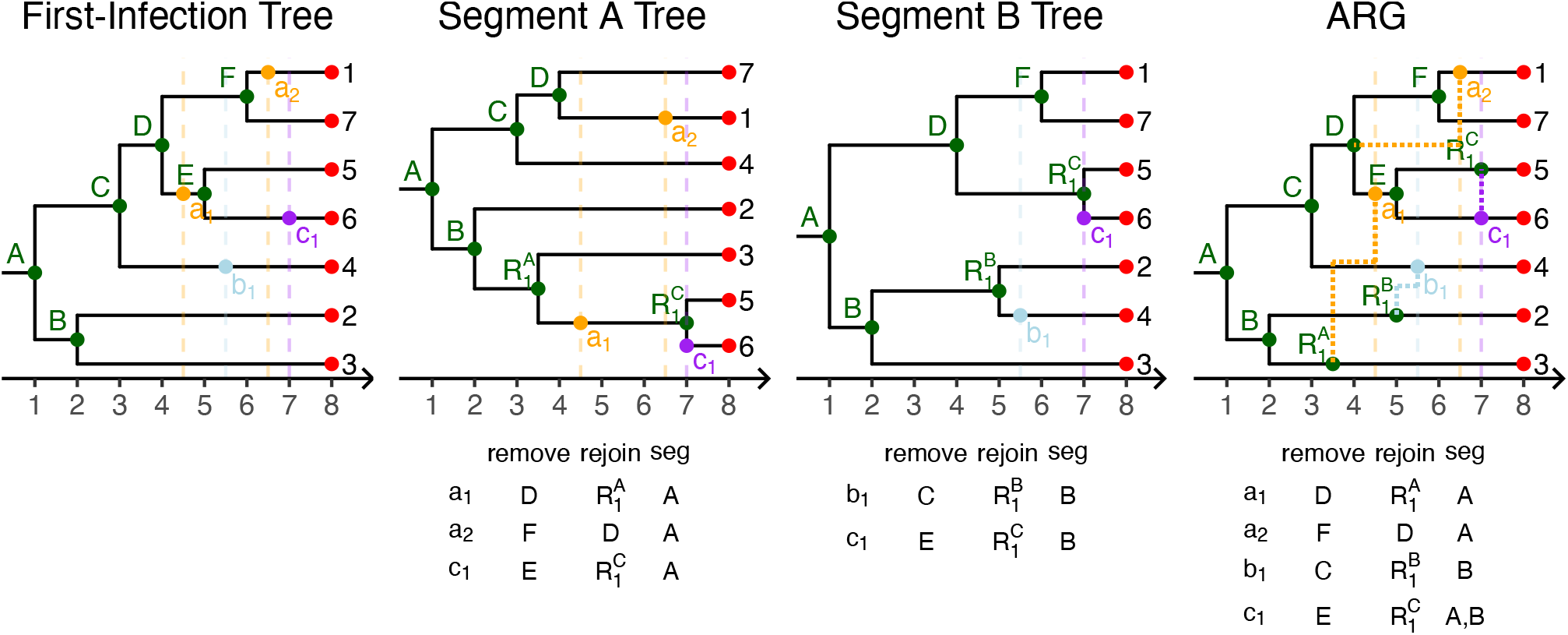
The number of visible reassortments and the remove-and-rejoin tables. By comparing the first-infection tree and each segment tree after dismissal, one can derive a remove-and-rejoin table for each segment tree. The union of all remove-rejoin tables reflects the true ARG and the number of visible reassortments. Nodes *a*_1_ and *a*_2_ indicate the location (*i*.*e*., the corresponding lineage and the reassortment time) of reassortments in segment A tree, *b*_1_ in segment B tree, and *c*_1_ in both.

**Figure S4:**
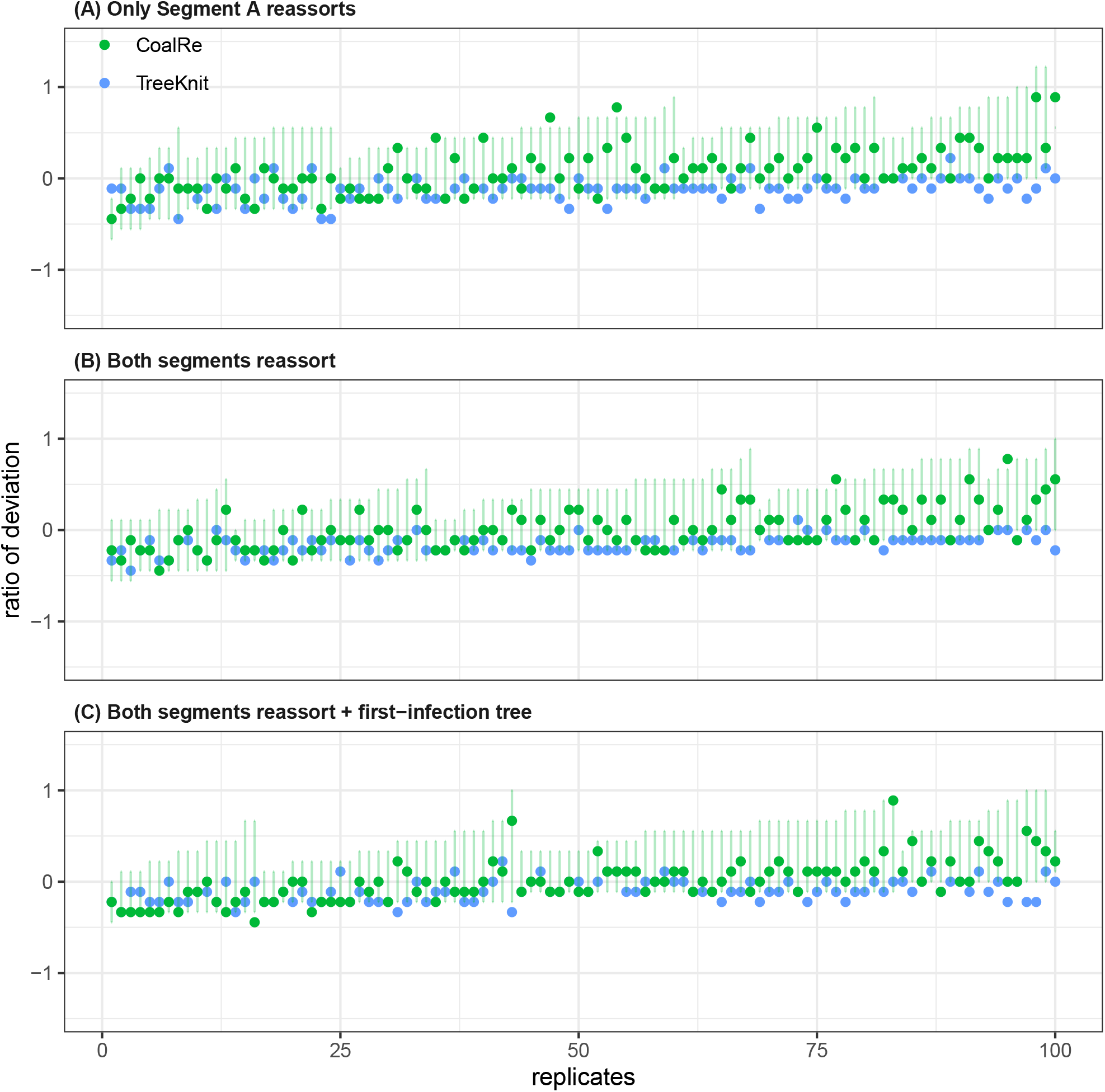
Closer look at the estimated number of reassortments by. `CoalRe` **and** `TreeKnit`. With a constant expected population size *I* = 100 and rates *λ* = 5, *µ* = 4.75, *Ψ* = .25, and *ρ* = .2, we simulated 100 sets of trees with 50 samples and 9 visible reassortment events. In `CoalRe`, 500 posterior ARGs are used to compute the maximum clade credibility tree (green dot) for each replicate, and the green bars indicate the 95% HPD among those realizations. Two reassortment schemes are discussed: in (A) only segment A reassorts, *ρ*_*A*_ = .2 and *ρ*_*B*_ = *ρ*_*AB*_ = 0; and in (B) and (C), both segments reassort but not simultaneously, *ρ*_*A*_ = .12, *ρ*_*B*_ = .08, and *ρ*_*AB*_ = 0. Two inference schemes are discussed: in (A) and (B), only segment trees are used for inference; in (C), segment trees and the simulated first-infection tree are used.

**Figure S5:**
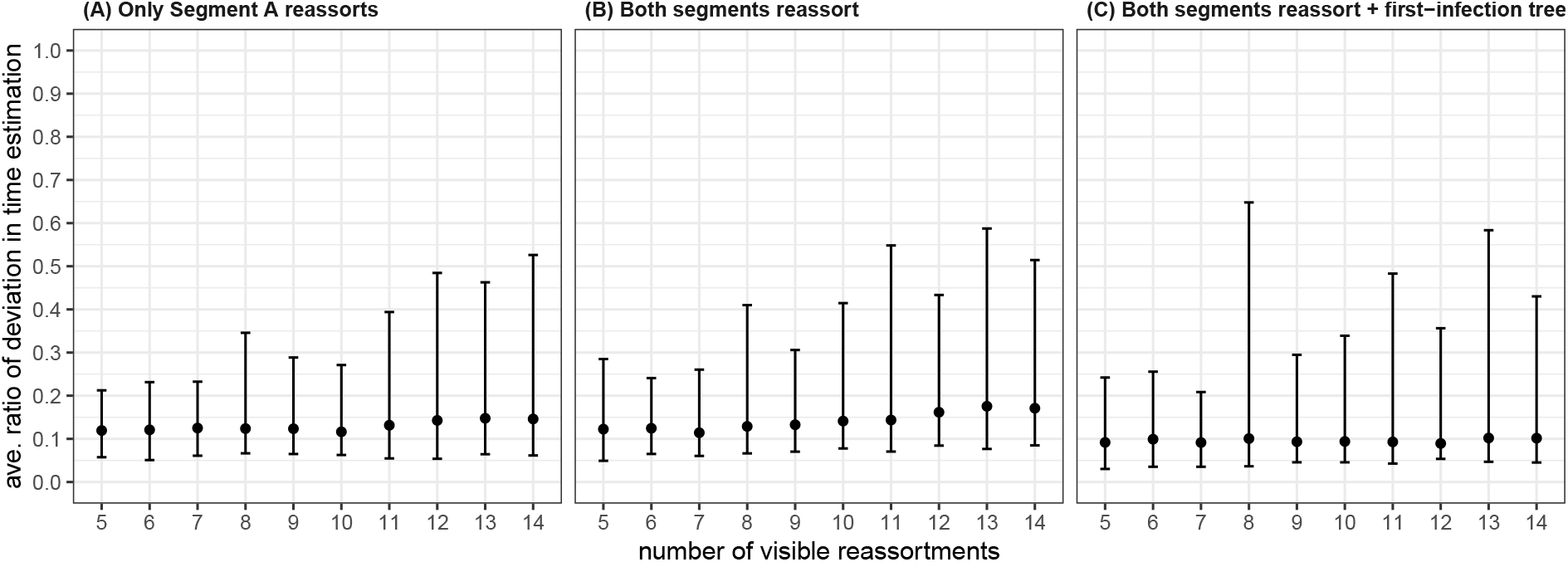
Deviation in reassortment time estimations by. `CoalRe`. With a constant expected population size *I* = 100 and rates *λ* = 5, *µ* = 4.75, *Ψ* = .25, and *ρ* = .2, we simulated 100 trees with 50 samples for each of the number of visible reassortments, ranging from 5 to 14. For each simulated tree, `CoalRe` produced 500 posterior ARGs and some of them estimated the number of visible reassortments correctly. Among those getting the correct estimation, we computed the average RMSE of reassortment time between each of the posterior ARG by `CoalRe` and the simulated tree, normalized by the height of the simulated tree. We define this metric as the *average ratio of deviation in time estimation*. We show the 95% quantile and the median of this metric for each of the number of visible reassortments. Two reassortment schemes are discussed: (A) only segment A reassorts, *ρ*_*A*_ = .2 and *ρ*_*B*_ = *ρ*_*AB*_ = 0, and (B) both segments reassort but not simultaneously, *ρ*_*A*_ = .12, *ρ*_*B*_ = .08, and *ρ*_*AB*_ = 0. Two inference schemes are discussed: in (A) and (B), only segment trees are used for inference; in (C), segment trees and the simulated first-infection tree are used.

## Notes

### Competing Interest Statement

The authors have declared no competing interest.

### Summary of Updates

Manuscript reorganized for clarify; Fig. 8, S5, and S6 revised to include the impact of the simulated first-infection tree on reassortment inference; Supplementary Material updated.

